# Dissecting the Network Architecture of a Plant Circadian Clock Model: Identifying Key Regulatory Mechanisms and Essential Interactions

**DOI:** 10.64898/2026.03.15.711848

**Authors:** Shashank Kumar Singh, Ashutosh Srivastava

## Abstract

Circadian rhythms are self-sustained biological oscillations that coordinate diverse physiological processes in plants, including growth, metabolism, and environmental responses. These rhythms arise from an interconnected transcriptional translational feedback network that integrates multiple entrainment cues such as light and temperature. The plant circadian clock is organized around key regulatory loops involving CCA1, LHY, PRRs, TOC1, ELF4, LUX, and other transcriptional regulators, whose coordinated interactions ensure precise and robust oscillations. In this study, we developed an ordinary differential equation based mathematical model, building upon a previous framework to incorporate additional regulatory modules and transcriptional controls that better reflect experimentally observed behaviour. To elucidate the regulatory organization of this model, we performed a multi-layered computational analysis combining four complementary approaches: (i) period sensitivity analysis to quantify how parameter perturbations influence the system’s timing, (ii) phase portrait analysis to visualize dynamic interactions among key components, (iii) knockout analysis to identify parameters essential for sustained rhythmicity, and (iv) network impact analysis using composite weighted network indices to evaluate hierarchical control across the network. Together, these analyses reveal that transcriptional repression, protein degradation, and light-regulated synthesis form the dominant control mechanisms within the circadian system. The results highlight a hierarchical and robust network structure centred on the CCA1/LHY and PRRs feedback loop, with redundant modules ensuring stability under perturbations. Thus, this model provides an improved, biologically consistent framework for dissecting the dynamic architecture of the plant circadian clock and guiding future experimental validation.

## 1. Introduction

Plants, like other living organisms, possess an internal biological timing system known as the circadian clock. This clock generates ∼24-hour rhythms in gene expression, metabolism, and physiology, enabling alignment of internal biological processes with predictable environmental cues such as light, temperature, and photoperiod(Atamian & Harmer, 2016; Inoue et al., 2018). Circadian rhythms play a central role in regulating key plant functions, including photosynthesis, hormone signaling, growth, flowering, and stress adaptation. At the molecular level, the plant circadian oscillator comprises interconnected transcriptional translational feedback loops (TTFLs) that coordinate rhythmic gene expression across day and night. In *Arabidopsis thaliana*, the core loop involves the morning repressors CCA1 (CIRCADIAN CLOCK ASSOCIATED 1) and LHY (LATE ELONGATED HYPOCOTYL), the evening regulator TOC1 (TIMING OF CAB EXPRESSION 1), and sequentially expressed PRR9, PRR7, and PRR5, which mediate daytime repression(Nohales & Kay, 2016). The Evening Complex (ELF3-ELF4-LUX) further integrates environmental signals to fine-tune evening gene expression(Mizuno et al., 2014, 2015).

Light quality and photoperiod have a strong influence on circadian entrainment and various aspects of plant growth. Different wavelengths activate distinct photoreceptor pathways, with red and far-red signals perceived by phytochromes, which adjust oscillator speed and traits such as hypocotyl elongation and shade avoidance(Oakenfull & Davis, 2017). Blue-light sensing cryptochromes contribute to period shortening and support photoperiodic control of flowering, while combined red-blue input under white light provides a stronger entrainment signal than either wavelength individually(Chen et al., 2004). Additional receptors, including ZEITLUPE and UVR8, integrate blue and UV-B cues to regulate components such as TOC1, PRR9 and GI, linking spectral composition to changes in gene expression and growth. Variation in day length further modulates oscillator phase, amplitude, and gating, demonstrating that both wavelength and photoperiod act together to coordinate circadian rhythms with developmental outputs(Oakenfull & Davis, 2017).

Mathematical models have been a critical tool in understanding this intricate regulatory network and its role in several crucial aspects of plant physiology including yield, flowering time, etc(Chan et al., 2024; Chew et al., 2022; Fogelmark & Troein, 2014; Foo et al., 2016; Locke, Millar, et al., 2005; Locke, Southern, et al., 2005; Pokhilko et al., 2010; Singh & Srivastava, 2024; Zeilinger et al., 2006). The early models described interactions between CCA1/LHY and TOC1 and successfully captured their basic oscillatory behavior(Locke, Millar, et al., 2005; Locke, Southern, et al., 2005). Subsequent models with interconnected regulatory loops integrated additional PRR components and post-translational regulators, improving simulation of light entrainment and mutant phenotypes(Pokhilko et al., 2010; Zeilinger et al., 2006). More recent models have incorporated the Evening Complex and regulators such as RVE8 and NOX, linking circadian control to broader physiological outputs including flowering time and hypocotyl elongation under variable environmental conditions(Chan et al., 2024; Chew et al., 2022; Fogelmark & Troein, 2014).

Despite these advances, existing models still face limitations in reproducing as well as predicting observed dynamics of the plant’s circadian clock under natural conditions. For example, Pay (2022) model was primarily developed to simulate hypocotyl growth under different light wavelengths (Red, Blue and Red+Blue) and durations (6L:18D, 4L:20D, and 2L:22D) by linking circadian regulation to photomorphogenic signaling(Pay et al., 2022). While this framework successfully captured growth responses under varying light conditions, it did not focus on the temporal expression patterns of individual clock components. Fogelmark model has been able to capture the individual expression of the genes in response to varying photoperiods but failed for some mutant conditions(Fogelmark & Troein, 2014). Additionally, the effect of varying light intensity has not been modeled in sufficient detail, leaving gaps in understanding how changes in irradiance shape the amplitude, phase, and robustness of core clock gene expression.

To address these gaps, we used Pay model as the basis for development of a more comprehensive model that could capture the expression dynamics of core clock components under varying light intensity and photoperiods. To do so, we first predicted the expression pattern of key circadian genes based on Pay model under natural light conditions at spring equinox and compared them to curated expression profiles by Nagano et al., 2019. The expression of key circadian genes in *Arabidopsis* was predicted based on Pay model. We found amplitude and phase difference between predicted and experimentally curated expression profiles (Figure S1).

We then developed an extended mathematical model (hereafter referred to as M1 Model), that expands the Pay (2022) model by incorporating a GI/ZTL (GZ) module and additional light-dependent inhibitory interactions. M1 Model represents a more complete integration of transcriptional feedback processes governing rhythmicity. In the sections that follow, we describe the structure and validation of M1 Model. We further employ four complementary analyses-period sensitivity, phase portrait, knockout, and network impact to dissect its architecture and dynamic behavior. These approaches together reveal the hierarchical organization, robustness, and key regulatory mechanisms underlying circadian timing in *Arabidopsis thaliana*.

## 2. Results

### 2.1 M1 Model: Design and Validation

The Pay model, when simulated under a natural light conditions at spring equinox (see methods), showed phase and amplitude deviations between the model predicted expression of core components-CCA1/LHY, PRR7/9, PRR5/TOC1, and ELF4/LUX and curated expression profiles obtained from the Nagano et al., 2019 study (Nagano et al., 2019) (Figure S1). These discrepancies indicated missing regulatory interactions limiting its ability to reproduce experimentally consistent oscillatory behaviour.

To address the limitations of the Pay model, we incorporated regulatory components and interactions supported by recent experimental studies (Figure 1A). Central to these updates is the inclusion of a GI/ZTL (GZ), which represents the combined regulation by GIGANTEA and ZEITLUPE. This complex is known to control the stability of evening-phased proteins such as PRR5 and TOC1 and therefore acts as an important node linking environmental light cues to clock protein expression(Cha et al., 2017; Kiba et al., 2007; Más et al., 2003). Building on this module, we introduced the GI/ZTL-dependent promotion of PRR5 and TOC1 degradation, a key process that shapes the timing of the evening loop(Más et al., 2003). We also incorporated degradation of GZ by COP1, reflecting the photoperiod-dependent regulation of the GI-ZTL complex through COP1-mediated light signaling(Jang et al., 2015). Finally, inhibition of PIF by CRY was added to represent cryptochrome-driven repression of PIF-mediated growth responses, allowing the model to better link light input pathways to downstream physiological outputs(Ma et al., 2016). Collectively, these updates strengthen the coupling between transcriptional and translational layers.

**Figure 1.**
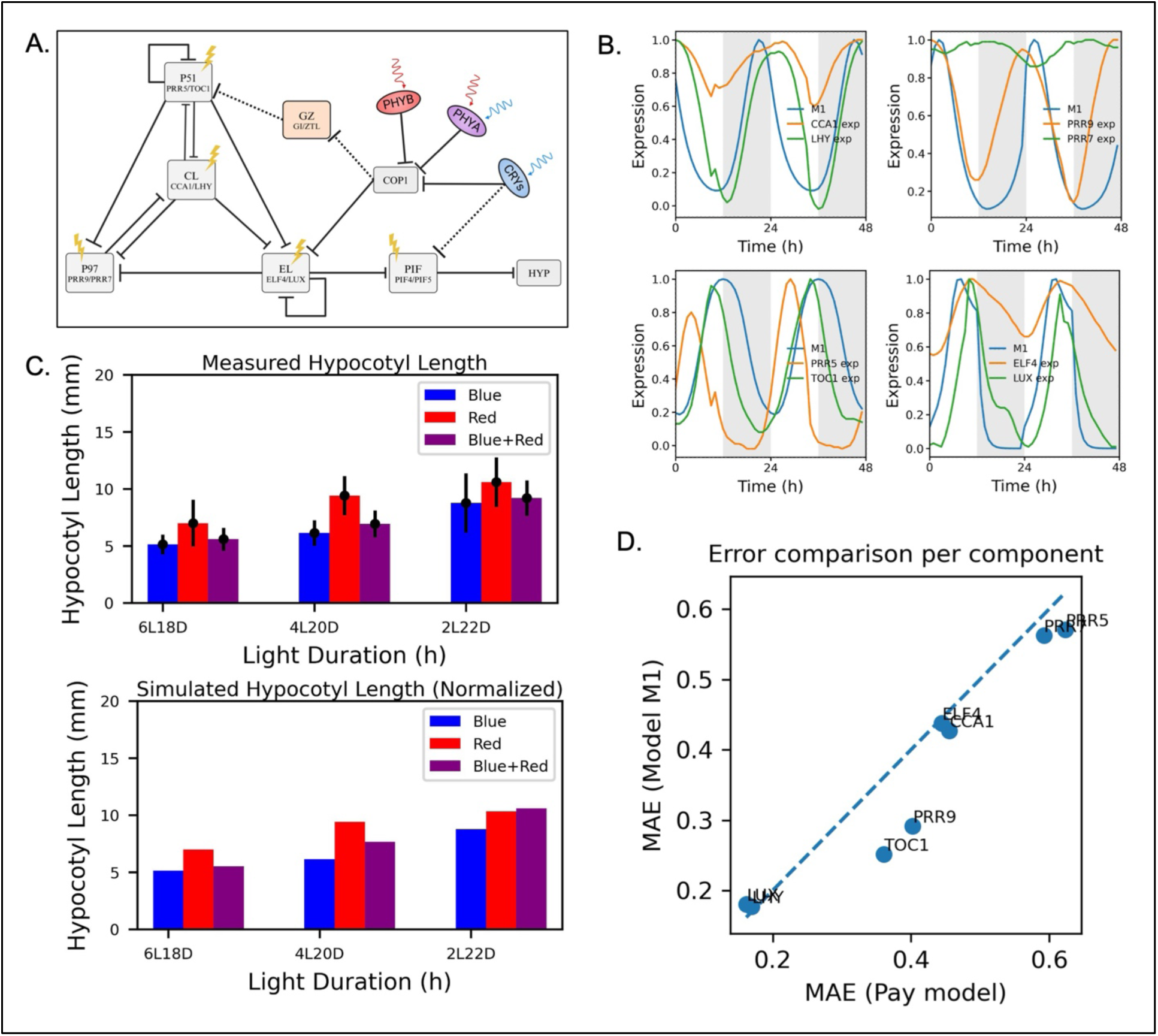
Model extension, validation, and predictive accuracy of hypocotyl growth under varying light conditions. **(A)** Schematic of the updated circadian regulatory network (M1 Model), adapted from Pay et al., 2022. The model incorporates GI and ZTL (shown in peach) and revises the inhibitory interactions from CRYs to PIF4/5, indicated by dashed lines. Solid arrows and T-bars denote activation and repression, respectively, within the network comprising core clock components (CL, P51, P97, EL), photoreceptors (PHYA, PHYB, CRY1/2), and growth regulators (COP1, PIFs, HYP). **(B)** Expression of core components of M1 model under natural condition at spring equinox with curated expression data extracted from the Nagano et al.,2019 study(Nagano et al., 2019). **(C)** Validation of model predictions through measured (left) and simulated (right) hypocotyl lengths under different light quality (blue, red, blue+red) and duration (6L18D, 4L20D, 2L22D) conditions. Error bars represent standard deviations in experimental measurements. **(D)** Mean Absolute Error (MAE) between experimental data and model predicted expressions of core components by M1 and Pay models under natural condition at spring equinox.

We first compared hypocotyl growth reported by Pay et al., 2022 with M1 model predictions across three light qualities (red, blue, and red+blue) and three photoperiods (6L:18D, 4L:20D, and 2L:22D) (Pay et al., 2022) (Figure 1C). The M1 model reproduced the experimentally observed growth patterns under all tested conditions (except for photoperiod of 2L22D under blue+red light), supporting the validity of the model.

Next, the M1 model was simulated under natural light conditions at equinoxes and solstices, and its predicted expression dynamics of CL (CCA1/LHY), P97 (PRR9/PRR7), P51 (PRR5/TOC1), and EL (ELF4/LUX) were compared with available curated expression profiles in natural conditions. The M1 model showed slightly better correspondence to the curated expression values in similar conditions as compared to the Pay model (Figure 1B). Further, comparison of Mean Absolute Error (MAE) between experimental expression profiles and model predictions showed consistently lower error for the M1 model relative to the Pay model, indicating improved predictive accuracy (Figure 1D;S2). Together, these results show that M1 replicates both molecular expression patterns and growth responses across diverse natural light regimes better than the Pay model.

Although the M1 model provides lower error with respect to the experimental data, there are considerable differences, highlighting the challenges in modeling the responses in natural conditions. To understand the contribution of different components to the overall stability and robustness of the model, we performed in silico parameter knock out, period sensitivity, and phase portrait analysis.

### 2.2 *In silico* knockout analysis identifies a compact core oscillator anchored by CL-P97-P51-EL interactions

To determine which components of M1 model are essential for sustaining circadian oscillations in *Arabidopsis*, we simulated the M1 Model with each parameter independently set to zero (virtual knockout) while keeping all others unchanged (see Methods) in two different conditions-24L0D and 12L12D (henceforth, LL and LD). The effect of knockout was measured as average change in the period of oscillation for four core clock components CL (CCA1/LHY), P97 (PRR9/PRR7), P51 (PRR5/TOC1), and EL (ELF4/LUX). Representative expression profiles illustrate two distinct responses to parameter knockout (Figure 2A). Removal of core regulatory parameters abolishes oscillations in CL, P97, P51, and EL, indicating loss of rhythmicity, whereas knockout of other parameters preserves sustained oscillations with only minor amplitude changes. These contrasting dynamics suggest that a small set of interactions form the rhythm-generating core of the oscillator, while other processes primarily modulate clock behaviour.

**Figure 2.**
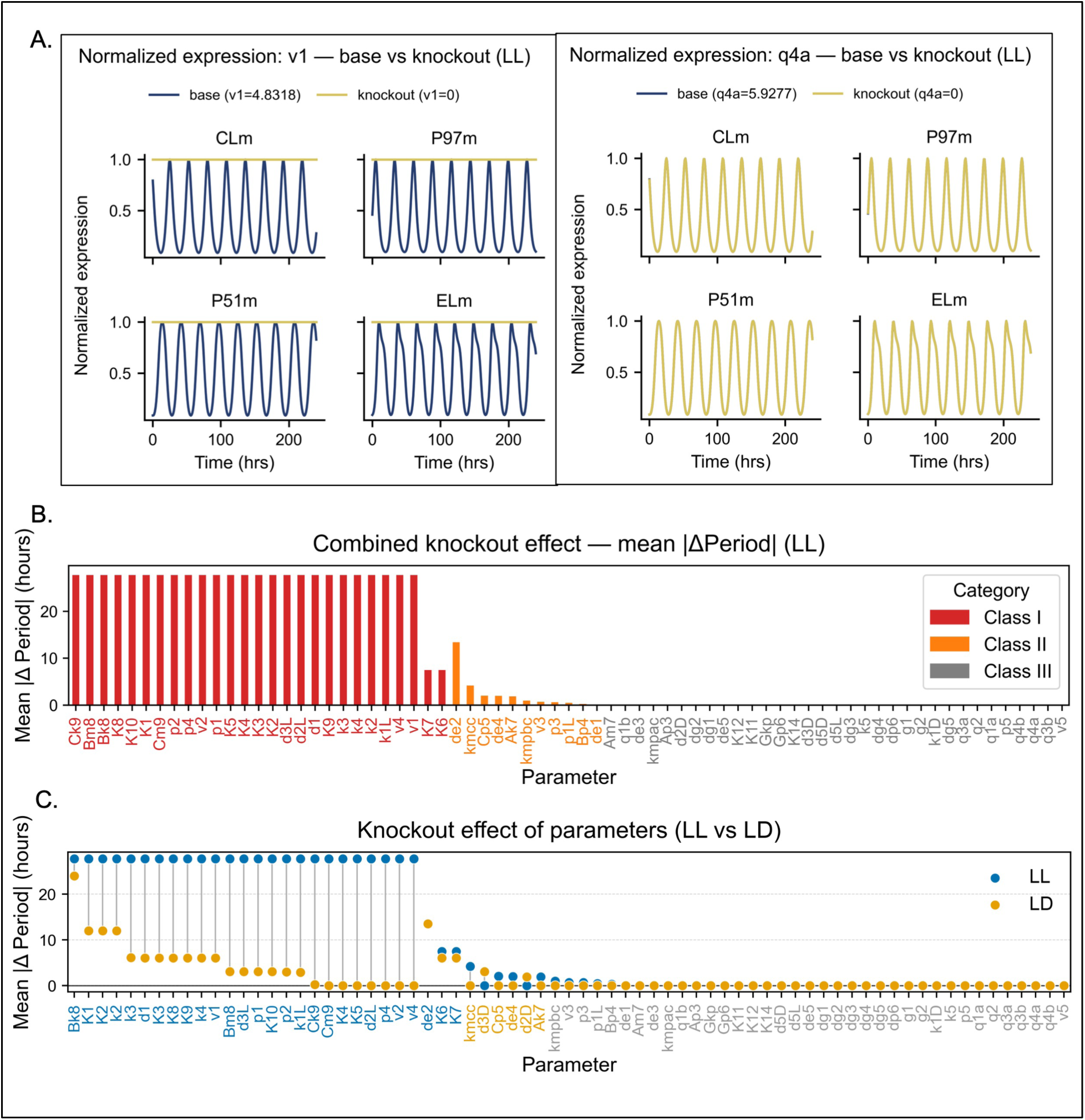
Combined knockout sensitivity of model parameters based on absolute period change. ***(A)*** Normalized expression profiles of the four core components (CLm, P97m, P51m, and ELm) under representative parameter variation. Left: variation of v1 (CL mRNA synthesis). Right: variation of q4a (light-regulated synthesis term). Blue and orange curves indicate reference and perturbed values, respectively. ***(B)*** Combined knockout effect of all parameters, quantified as mean absolute change in period (hours). Parameters are grouped into three classes: Class I (red, strong period disruption), Class II (orange, moderate effect), and Class III (grey, minimal effect). ***(C)*** Comparison of knockout-induced period changes under constant light (LL, blue) and light-dark cycles (LD, yellow). Several parameters show strong effects in LL but reduced impact in LD, indicating partial buffering by environmental entrainment.

The average change in period ranged from 0 to 27.7 hours as compared to the original model. We classified the parameters into three broad classes based on the effect of their knockout on average period. First (hereafter referred to as Class I), where the knockout of the parameter led to loss of rhythmicity, second (Class II), where the effect on period was moderate and third (Class III), where there was no effect on the period (Figure 2B).

Class I parameters abolished oscillations when knocked out, which shows that they form the rhythm-generating core of the model. These included key production and degradation steps that control CCA1/LHY (CL), PRR9/PRR7 (P97), PRR5/TOC1 (P51), and the evening complex (EL) (Table S1). Essential processes were CL synthesis (v1), translation (p1) and degradation (d1, k1L), P97 synthesis (v2), translation (p2), and mRNA degradation (k2), as well as P51 mRNA degradation (k3). Light dependent protein turnover was also critical, including P97 degradation in light (d2L) and P51 degradation in light (d3L). Parameters regulating EL synthesis (v4), translation (p4) and mRNA degradation (k4) were also critical. Feedback repression is a critical component of circadian regulatory circuit(Li et al., 2025). Consistent with this we found mRNA inhibition of CL by P97 (K1) and by P51 (K2), inhibition of P97 by CL (K3), by P51 (K4), and by EL (K5), and repression within the evening and P51 modules, including inhibition of P51 by CL(K6) and its self-inhibition (K7), inhibition of EL by CL (K8), by P51 (K9) and its self-inhibition (K10) to be crucial.

Although, K6 and K7 have lower mean change in period than de2 they are classified in Class I due to complete loss of rhythmicity of P51 caused by their knockout. In addition, photoreceptor turnover contributed to oscillator integrity, including Michaelis-Menten constants governing PhyB (Bk8) and Cry (Ck9) degradation, PhyB degradation (Bm8) and Cry degradation (Cm9). Together, these parameters define the tightly coupled CL-P97-P51-EL network, supported by regulated photoreceptor decay, that sustains circadian oscillations. This structure is consistent with experimental evidence placing the CCA1/LHY-PRR9/7 repression loop at the core of the *Arabidopsis* clock(Green et al., 2002; Más et al., 2000).

Class II parameters altered period but preserved rhythmicity, which identifies them as timing regulators rather than rhythm generators. These included translational and synthesis steps that adjust the strength and delay of signal propagation through the network. Light-induced CL translation (p1L), P51 mRNA synthesis (v3) and translation (p3), EL degradation (de1), PhyB translation (Bp4) and Cry translation (Cp5) all modulated the pace of oscillations without abolishing them. Photoreceptor kinetics also shaped circadian timing. The binding rates of COP1 to Cry (kmcc) and PhyB (kmpbc), the Michaelis-Menten constant governing PhyA degradation (Ak7), COP1-mediated EL degradation (de2) and COP1-PhyB mediated EL degradation (de4) influence how strongly and how long light signals persisted in the system. These parameters likely adjust effective light sensitivity and signal decay, which in turn shifts circadian period. Together, these parameters form a flexible tuning layer that modulates circadian pace through translational control, photoreceptor turnover, and evening-loop repression strength, while preserving the underlying oscillatory core(Su et al., 2021).

Class III parameters had minimal impact on period, which indicates buffering within the extended regulatory network. Several light-input and degradation terms showed little effect on clock pace. These included CL light-induced synthesis through PhyA, PhyB, and Cry (q1a, q3a, q4a), and CL mRNA degradation in dark (k1D). Likewise, EL degradation through multiple routes COP1-PhyA mediated degradation (de3), and COP1-Cry mediated degradation (de5) produced only minor period shifts. Light-induced P97 synthesis through PhyA, PhyB, and Cry (q1b, q3b, q4b), as well as its light-independent degradation rate (q2) and dark degradation (d2D), were also weak modulators of timing. Similar buffering was observed for P51 degradation in dark (d3D), PhyA degradation (Am7), PhyA (Ap3), and COP1-PhyA binding (kmpac). Peripheral modules downstream of the core oscillator remained largely insulated. PIF turnover and synthesis terms including PIF mRNA degradation (k5), PIF translation and synthesis (p5, v5), PIF protein degradation in dark and light (d5D, d5L) has no influence on period. The entire GZ module - translation (Gp6), degradation via multiple routes (dg1, dg2, dg3, dg4, dg5), P51 degradation by GZ (dp6) and Michaelis-Menten constant (Gkp) was similarly insensitive to knockout. Regulatory links such as inhibition of PIF by EL (K11) and Cry (K14), activation of hypocotyl growth by PIF (K12), and the dissociation rate (kd) also showed minimal timing effects. Downstream outputs, including baseline and PIF-induced hypocotyl growth (g1, g2), were similarly buffered. These results reflect biological robustness in the plant clock. Light input, photoreceptor turnover, and growth output pathways are distributed across overlapping regulatory layers, which seem to absorb perturbations without disrupting core circadian timing.

Under LD conditions, the overall hierarchy of the M1 model parameters described above remains consistent, but the distribution of essential parameters shifts compared to constant light (LL) (Figure 2C). In LL, the oscillator depends strongly on core transcriptional repression (CL-P97-P51-EL) and regulated light-dependent degradation, because light input is continuous and internal feedback must sustain rhythmicity. In LD, external day-night entrainment partially stabilizes oscillations, and several parameters that were essential in LL move to Class II or Class III. Parameters that shift from Class I in LL to Class II in LD include CL translation (p1), CL mRNA degradation (k1L), P97 translation (p2), PhyB degradation (Bm8), P51 degradation in light (d3L), EL self-inhibition (K10), and Michaelis-Menten constant of Cry degradation (Ck9). Parameters that lose essentiality entirely under LD condition shifting to Class III include P97 synthesis (v2), P97 light-dependent degradation (d2L), EL synthesis (v4), EL translation (p4), Cry degradation rate (Cm9), and inhibition of P97 by P51 (K4) and by EL (K5). Notably, the conserved Class I core under LD comprises of CL synthesis (v1) and degradation (d1), P97 mRNA degradation (k2), P51 mRNA degradation (k3), EL mRNA degradation (k4), the Michaelis-Menten constant for PhyB degradation (Bk8), and the inhibitions K1, K2, K3, K6, K7, K8, and K9. Additionally, dark-phase degradation of P97 (d2D) and P51 (d3D) that are Class III in LL gain period-modulation roles in LD likely because dark phase imposes a distinct degradation window absent in constant light. COP1-mediated EL degradation (de2) is Class II in both LL and LD conditions, indicating its stable role of timing regulator being independent of photoperiod.

Together, this contrast shows that under constant light the clock behaves as a self-sustained autonomous oscillator dominated by core repression loops, whereas under LD it functions as a driven system where light-dark transitions redistribute control. The LD oscillator operates with a reduced but sufficient Class I core, centred on mRNA turnover of CL, P97, P51 and EL, CL synthesis and degradation, PhyB degradation and seven inhibitory interactions (K1, K2, K3, K6, K7, K8 and K9). While light-dependent protein degradation of P97 (d2L), EL synthesis (v4), EL translation (p4) and additional inhibitory interactions (K4 and K5) become dispensable. This shift supports the idea that environmental cycles buffer some core interactions while increasing dependence on modules that mediate entrainment and phase coordination.

Collectively, the knockout analysis shows that M1 Model captures a clear hierarchical structure: a core set of essential feedback and degradation processes, a flexible layer that tunes period, and a redundant layer that ensures robustness. This organisation mirrors experimental observations from *Arabidopsis*, highlighting the interplay between core transcriptional loops, light-dependent modulation, and buffered peripheral pathways. After identifying which parameters are essential, modulatory, or redundant, we next examined how variation in each parameter value influences circadian timing using period sensitivity analysis.

### 2.3 Period sensitivity analysis reveals that timing is controlled by a few high-impact transcriptional and light-dependent processes

To quantify how parameter variation influences circadian timing, each parameter was varied across its defined range while monitoring period changes in the four core clock components. Representative expression profiles illustrate three distinct responses of the oscillator to parameter variation (Figure 3A). Changes in certain parameters shorten the oscillation period, while others extend the period, as reflected by shifts in the spacing of successive peaks of CL, P97, P51, and EL. In contrast, variation in some parameters produces little or no change in oscillatory timing. These contrasting behaviours indicate that circadian period in the M1 model is controlled by a limited set of highly sensitive regulatory processes, whereas many parameters have minimal influence on clock timing. The analysis revealed three distinct sensitivity groups: highly sensitive, moderately sensitive, and insensitive parameters (Figure 3B).

**Figure 3.**
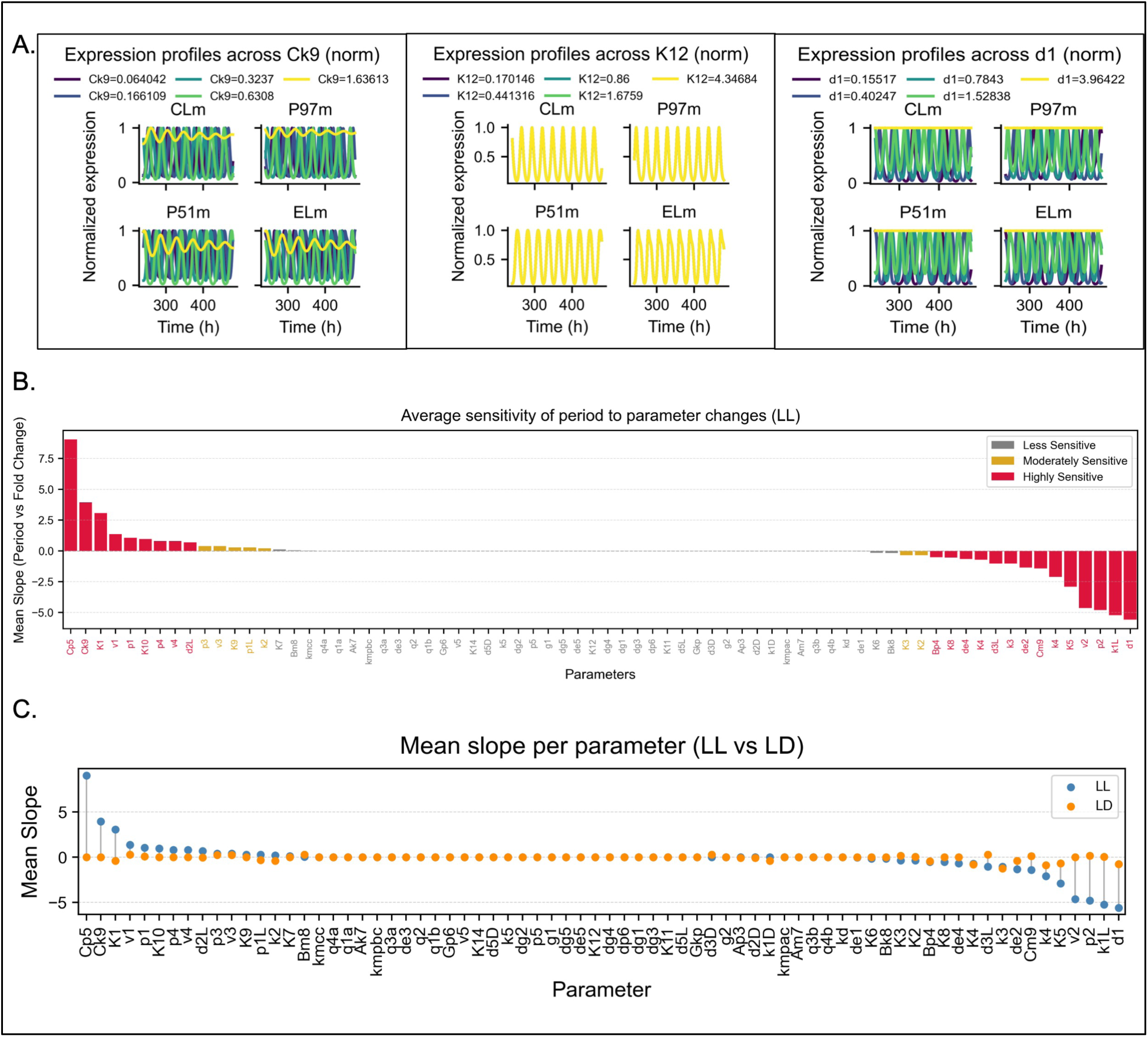
Average sensitivity of circadian period to fold change in model parameters. ***(A)*** Normalized expression profiles of the four core components (CLm, P97m, P51m, and ELm) across graded variation of representative parameters (Ck9, K12, and d1). Multiple colored traces indicate increasing parameter values. Changes in repression strength alter oscillation amplitude, phase spacing, and waveform sharpness while preserving sustained rhythmicity over most ranges. ***(B)*** Average sensitivity of period to parameter changes under constant light (LL), quantified as the mean slope of period versus fold change. Positive slopes indicate period lengthening with parameter increase, while negative slopes indicate period shortening. Parameters with the strongest absolute slopes cluster in core synthesis, degradation, and repression terms, confirming their dominant role in tuning oscillator pace. ***(C)*** Dumbbell comparison of mean period slopes under LL (blue) and light-dark cycles (LD, orange). Several parameters show strong sensitivity in LL but reduced slopes in LD, indicating environmental buffering under entrainment.

Highly sensitive parameters caused strong changes in period, which shows that these processes tightly constrain circadian timing. This group included core production and degradation steps of the morning loop. CL synthesis (v1), translation (p1), degradation (d1), and light-dependent mRNA degradation (k1L) were particularly sensitive, indicating that precise control of CCA1/LHY levels is essential for stable timing. P97 synthesis (v2), translation (p2), and light-dependent degradation (d2L) were also highly sensitive, which highlights the importance of balanced PRR9/7 turnover in setting period length. Key inhibitory interactions within the core loop further shaped timing. Inhibition of CL by P97 (K1) and inhibition of P97 by EL (K5) showed strong effects on period, reflecting the tight coupling between the morning and evening modules. Small changes in these repression strengths led to measurable shifts in period (Figure 3A, B). Inhibition of P97 by P51 (K4) and inhibition of EL by CL (K8) were also highly sensitive, further linking repression dynamics across all three loops. Photoreceptor-linked processes also constrained clock pace. The Michaelis-Menten constant of Cry degradation (Ck9) and Cry translation (Cp5) significantly altered the period, which suggests that Cry stability and abundance modulate light signal persistence and feedback strength. PhyB translation (Bp4) likewise showed high sensitivity, indicating that photoreceptor abundance constrains clock pace under constant light. EL synthesis (v4) and translation (p4) were highly sensitive, underscoring the importance of evening-loop protein accumulation in setting period length. COP1-mediated EL degradation (de2) and PhyB-COP1 mediated degradation of EL (de4) were also highly sensitive, indicating that EL turnover routes through COP1 strongly constrain timing. P51 mRNA degradation (k3), EL mRNA degradation (k4), and P51 light-dependent degradation (d3L) were similarly highly sensitive, reflecting the contribution of post-transcriptional regulation to period control. EL self-inhibition (K10) also fell in the highly sensitive group, consistent with strong self-regulatory control of the evening loop. Together, these results show that the quantitative balance between CL-P97 mutual repression, light-dependent CL turnover, P97 stability in light, P51 and EL turnover, evening-loop synthesis and self-regulation, COP1-mediated EL degradation, and Cry kinetics are major determinants of the circadian period(Alabadí et al., 2002; Mizoguchi et al., 2002; Nakamichi et al., 2005, 2010; Wang et al., 2021).

Moderately sensitive parameters shifted period with smaller magnitude, which indicates that they fine-tune timing without controlling it. This group included core expression steps such as P97 mRNA degradation (k2), P51 mRNA synthesis (v3) and translation (p3). Inhibition of CL by P51 (K2), P97 by CL (K3), and EL by P51 (K9) were also moderately sensitive, indicating that repression interactions in the flanking loops provide additional period tuning. These processes adjust protein accumulation rates and phase relationships but do not disrupt oscillatory timing. Light-linked regulation also showed moderate influence. CL light-induced translation (p1L) altered period modestly, which suggests that photoreceptor-coupled translational control tunes light sensitivity rather than set clock speed directly(Devlin & Kay, 2000). Regulatory feedback within the evening module contributed additional modulation. Together, these parameters act as secondary tuning nodes and interactions. They adjust translation efficiency, repression thresholds, and morning-loop feedback strength, thereby refining circadian pace while leaving the core timing architecture intact.

Insensitive parameters produced negligible changes in period across their tested ranges. Many dark-dependent and peripheral degradation processes fell into this group, including P97 degradation in dark (d2D), P51 degradation in dark (d3D) and PIF degradation in dark and light (d5D, d5L). Multiple EL degradation routes basal turnover (de1), and COP1 complexes with PhyA and Cry (de3, de5) also showed minimal influence on timing. Photoreceptor-associated kinetics were largely buffered. These included the Michaelis-Menten constants and degradation rates of PhyA and PhyB (Ak7, Am7, Bk8, Bm8), PhyA translation (Ap3), and COP1 binding rates to Cry, PhyA, and PhyB (kmcc, kmpac, kmpbc). Light-induced synthesis terms for CL and P97 through PhyA, PhyB, and Cry (q1a, q1b, q3a, q3b, q4a, q4b), as well as the light-independent degradation constant (q2), likewise had little effect on period. Inhibition terms within the core loop like P51 by CL (K6) and P51 by itself (K7) within this category show higher sensitivity than other regulatory interactions outside the core loop. These included inhibition terms such as PIF by EL (K11), and PIF by Cry (K14), along with the dissociation rate (kd). mRNA degradation terms for CL in dark (k1D), and PIF (k5) were also weak contributors.

Downstream output and peripheral synthesis processes, including PIF translation and synthesis (p5, v5), and hypocotyl growth parameters (g1, g2, K12), showed minimal timing impact. Together, these results show that most light-input, degradation, and growth-output pathways are distributed across redundant regulatory layers. This buffering preserves circadian period while allowing flexibility in downstream responses, which supports overall system robustness.

Under LD condition, period sensitivity shifts compared to LL (Figure 3C). In LL, circadian timing is tightly constrained by core CL-P97 repression, light-dependent CL turnover, P97 stability in light, P51 and EL turnover, evening-loop synthesis and self-regulation, COP1-mediated EL degradation, and Cry kinetics. In LD, most of these core parameters, including CL synthesis (v1), CL translation (p1), P97 translation (p2), P97 synthesis (v2), EL synthesis (v4), EL translation (p4), light-dependent degradation terms (k1L, d2L, d3L, de2, de4), and repression terms such as K1 and K8, become less or moderately sensitive, even though several remain structurally essential in knockout analysis. Instead, only a few parameters show high sensitivity, notably P51 mRNA degradation (k3), EL mRNA degradation (k4), CL degradation(d1), and inhibition of P97 by P51 and by EL (K4 and K5), highlighting a stronger role for mRNA turnover and core loop repression under cycling light. Comparison of knockout classification and quantitative period sensitivity reveals a clear separation between structural necessity and kinetic control within the clock. Parameters that are both Class I and highly sensitive in LD condition are P51 mRNA degradation (k3), EL mRNA degradation (k4), and CL degradation(d1). They sustain oscillations and strongly constrain the period under entrainment. Inhibition of P97 by P51 and by EL (K4 and K5) are highly sensitive in LD but are Class III in knockout analysis, indicating that their kinetic strength determines period without being individually required for rhythmicity. Conversely several Class I parameters in LD (K2, K3, K6-K9) sustain oscillations but show negligible period sensitivity, confirming structural but not kinetic essentiality. In contrast, several parameters are essential for network integrity but weak in period tuning under LD. Bm8, de2 and d3L are Class II in LD but with moderate period sensitivity, while Cm9 is Class III with negligible period effect. Others, including p1 and p2 (Class II in LD), modulate period without abolishing rhythms. Meanwhile, most dark degradation processes, PIF/GZ modules, growth outputs (g1, g2), and many light-induced transcription terms show negligible timing effects. Together, this layered structure explains how the plant clock maintains robustness through a tightly constrained core while allowing flexible tuning through peripheral regulatory modules.

With the key period determining parameters identified, we next examined how these regulatory processes shape system dynamics using phase portrait analysis to visualize oscillatory stability, amplitude, and coupling behaviour.

### 2.4 Core feedback maintains oscillatory structure while light inputs shift phase relationships

To examine how parameter variation alters the dynamic behaviour of the M1 model beyond period alone, we quantified changes in phase portrait geometry using the slopes of mean delta area and mean delta eccentricity. Phase portraits of pairwise combinations among the core components CL, P97, P51, and EL reveal three distinct dynamic responses to parameter variation (Figure 4A). Changes in some parameters lead to expansion or contraction of the limit-cycle trajectory, reflecting alterations in oscillation amplitude, whereas others modify the curvature and shape of the trajectory, indicating shifts in phase relationships between components. In contrast, several parameters produce minimal changes in phase-space geometry. To quantify these effects beyond qualitative observation, we measured changes in phase portrait geometry using the slopes of mean delta area and mean delta eccentricity. Phase portrait area reflects the overall amplitude of oscillations, representing the extent of coordinated molecular activity across the cycle, while eccentricity captures trajectory shape and symmetry, indicating how evenly different phases progress. The slopes of these metrics therefore quantify how strongly each parameter alters oscillation strength and waveform structure across its variation range.

**Figure 4.**
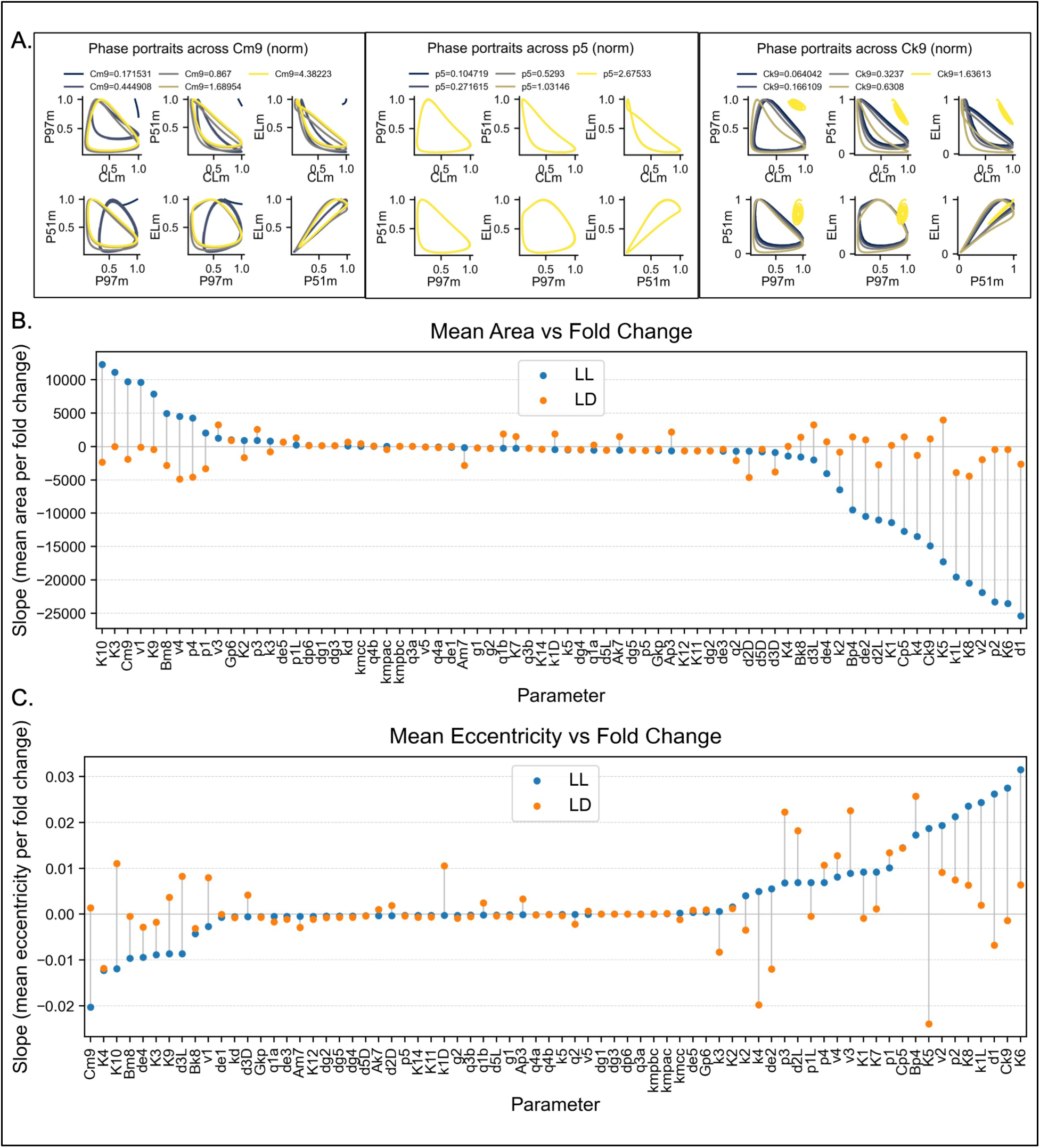
Phase portrait sensitivity and geometric reshaping of oscillations in the M1 model. ***(A)*** Phase portraits of pairwise combinations among the four core components (CLm, P97m, P51m, and ELm) across graded variation of representative parameters (Cm9, p5, and Ck9). Each colored trajectory corresponds to a different fold change. Changes in parameter strength alter limit-cycle size, curvature, and loop coupling. ***(B)*** Dumbbell comparison of mean slope of phase-space area versus fold change under constant light (LL, blue) and light-dark cycles (LD, orange). Positive slopes indicate expansion of the limit cycle, while negative slopes indicate contraction. ***(C)*** Mean slope of eccentricity versus fold change under LL and LD. Eccentricity captures trajectory shape and asymmetry rather than size. Most parameters cause modest shape changes, but selected synthesis and degradation terms induce stronger waveform distortion, particularly in LL.

Parameters that showed the largest area slopes were concentrated in the regulatory core, indicating strong control over oscillatory amplitude (Figure 4B). CL degradation (d1) produced the strongest area change, followed by inhibition of CL by P51 (K6), P97 translation (p2), P97 synthesis (v2), inhibition of EL by CL (K8), light-dependent CL mRNA degradation (k1L), inhibition of P97 by EL (K5), and Cry degradation (Ck9). These parameters are largely Class I in knockout, and their steep area slopes indicate that core degradation, translational control, and mutual repression within the CL-P97-EL axis set the strength of rhythmic molecular accumulation and degradation throughout the day. Biologically, these processes govern how strongly morning and evening regulators suppress each other and how rapidly transcripts and proteins are turned over, thereby controlling the intensity of circadian outputs.

In contrast, eccentricity slopes were smaller in magnitude and more selective, reflecting effects on waveform shape rather than oscillation strength (Figure 4C). Inhibition of CL by P51 (K6), Cry degradation (Ck9), CL degradation (d1), light-dependent CL mRNA degradation (k1L), and inhibition of EL by CL (K8) showed the largest eccentricity changes, along with P97 translation (p2) and Cry degradation (Cm9). These parameters altered trajectory symmetry and phase progression without producing large amplitude shifts, indicating refinement of timing relationships and waveform structure rather than disruption of the core oscillatory mechanism. Notably, PhyB translation (Bp4) and Cry translation (Cp5) showed moderate eccentricity effects, suggesting a secondary contribution of photoreceptor kinetics to waveform shaping.

Most peripheral and redundant processes showed comparatively weak slopes. Dark degradation pathways, output growth terms, and many light-induced transcription steps produced limited changes in both area and eccentricity, and the limit cycle remained closed and stable across their ranges. Together, these results demonstrate hierarchical control of dynamic structure: core degradation, translational steps, and mutual repression set oscillation amplitude, additional degradation and repression kinetics tune waveform shape, and peripheral branches contribute minimally to robust rhythmicity. This pattern mirrors the separation observed in knockout and period-sensitivity analyses and reinforces the central role of the CL-P97-P51-EL architecture in maintaining oscillatory integrity.

Under LD conditions, phase portrait analysis reveals a redistribution of geometric control compared to constant light. Area slopes now reflect how strongly light-driven processes modulate oscillatory amplitude under daily entrainment. The largest area slopes are dominated by EL synthesis (v4), P97 dark degradation (d2D), EL translation (p4), inhibition of EL by CL (K8), inhibition of P97 by EL (K5), followed by light-dependent CL mRNA degradation (k1L), P51 dark degradation (d3D), and CL translation (p1). This indicates that under cycling light, oscillatory expansion and contraction depend strongly on evening-module synthesis and degradation, core repression interactions, and CL translational control, which together determine how external light cues are translated into rhythmic molecular responses. PhyB translation (Bp4) and Cry translation (Cp5) show comparatively moderate area slopes, indicating a secondary rather than dominant role in amplitude control under LD.

Parameters with moderate area slopes under LD condition include P51 synthesis (v3), P51 light-dependent degradation (d3L), PhyB degradation (Bm8), and Cry degradation (Cm9). CL synthesis (v1) and repression terms K6 and K9 show very small area slopes under LD, indicating that their geometric influence on oscillatory amplitude is substantially reduced under entrainment compared to constant light. While these processes remain structurally essential for rhythm generation, their quantitative influence on oscillation amplitude is buffered by daily environmental entrainment.

Eccentricity slopes show a distinct pattern, reflecting control over waveform asymmetry and phase progression. The largest waveform distortions arise from PhyB translation (Bp4), inhibition of P97 by EL (K5), P51 synthesis (v3), P51 translation (p3) and EL mRNA degradation (k4), followed by light-dependent P97 degradation (d2L) and Cry translation (Cp5). These parameters modify phase balance and trajectory shape without proportional changes in amplitude, indicating regulation of timing coordination among network components. Notably, most dark degradation pathways, COP1 binding rates, growth outputs, and peripheral synthesis terms display minimal slopes in both metrics, and the limit cycle remains closed and stable across their ranges.

A further feature of the eccentricity analysis is the condition-dependent reversal of slope signs for several parameters when transitioning from LL to LD. Five parameters-d3L, k1D, Ap3, q1b, and Ak7, show concordant negative-to-positive shifts in both area and eccentricity slopes, indicating that under LD entrainment these processes switch from suppressing to amplifying both oscillation amplitude and waveform curvature, most likely because periodic light inputs activate their regulatory context and redirect their geometric influence on the limit cycle. Conversely, P51 mRNA degradation (k3) alone shows a concordant positive-to-negative shift in both metrics, implying that the destabilising effect of P51 transcript turnover under LL is reversed under LD, where light-driven P51 replenishment buffers the degradation signal. A subset of parameters displays discordant shifts, where area and eccentricity slopes reverse in opposite directions. Inhibition of P97 by EL (K5), Michaelis constant for Cry degradation (Ck9), inhibition of CL by P97 (K1), and EL degradation via COP1 (de2) gain amplitude-amplifying influence while simultaneously losing their waveform-expanding effect, suggesting that their degradation and repression kinetics are differentially channelled into amplitude versus phase-relationship control under entrainment. The complementary discordant pattern is seen in Cm9, K10, K9, and v1, which lose amplitude-expanding influence under LD while acquiring increased capacity to distort phase portrait shape. Together, these reversals demonstrate that light-dark cycling does not merely scale but fundamentally redirects the geometric roles of key regulatory parameters, reflecting a condition-dependent reorganisation of how amplitude control and waveform shaping are partitioned across the network.

Together, these results show that under LD conditions, geometric control shifts toward parameters that regulate evening-module synthesis and turnover, core repression interactions, and CL translational steps. PhyB translation and EL-linked repression dominate waveform shaping through eccentricity, while EL synthesis, P97 dark degradation, and core inhibition terms dominate oscillation amplitude through area, whereas many intrinsic core degradation and repression steps exert weaker geometric influence than in constant light. This pattern supports a driven-oscillator framework in which environmental cycling redistributes dynamic control from intrinsic core kinetics toward light-mediated entrainment modules while preserving overall oscillatory stability.

To place these effects within the broader regulatory architecture, we next integrated all metrics into a unified network-level framework.

### 2.5 The *Arabidopsis* clock operates through a tight core oscillator supported by adaptive light and stability modules

To integrate structural and dynamical contributions of each regulatory interaction into a single quantitative framework we performed network impact analysis. Under constant light (LL), we constructed a composite weighted, directed gene–gene network by combining knockout class, normalized knockout-induced period change, period sensitivity, and phase portrait metrics (area and eccentricity changes). For each interaction, these normalized terms were averaged to obtain a composite edge weight, which reflects structural necessity, control of oscillation pace, and influence on waveform geometry. Node influence was then assessed using weighted connectivity measures, allowing identification of dominant regulatory hubs and interaction pathways within the integrated network.

Integrating knockout, period sensitivity, phase portrait, and network impact analyses under constant light (LL) reveals a coherent and hierarchical regulatory architecture within the M1 model (Figure 5A). The composite LL network highlights CL and P97 as the dominant hubs, with the strongest weighted interactions concentrated within the CL–P97–P51–EL module. These nodes display the highest composite influence scores, reflecting their combined structural necessity, quantitative control of period, and impact on oscillatory geometry. In particular, CL degradation (d1), P97 translation (p2), light-dependent CL mRNA degradation (k1L), and P97 synthesis (v2) emerge as the highest weighted individual interactions, followed by Cry degradation (Ck9) and inhibition of EL by CL (K8), P97 by EL (K5) and CL by P97 (K1). The network topology therefore reinforces conclusions from knockout and sensitivity analyses that the CL–P97 feedback loop forms the quantitative backbone of the clock under continuous light.

**Figure 5.**
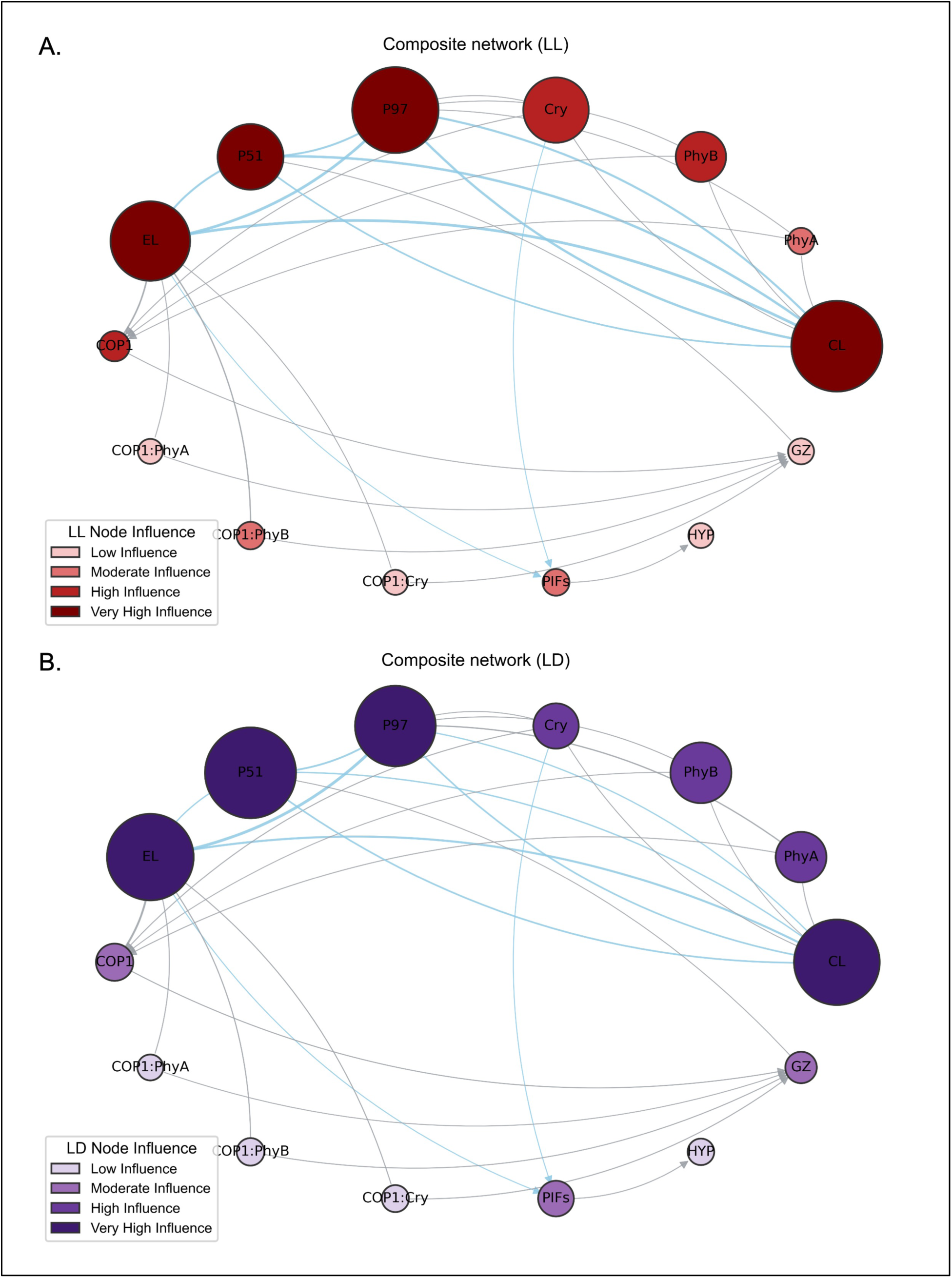
Composite directed network based on the influence of parameters and components on each other under **(A)** constant light (LL) and **(B)** light-dark cycle (LD). Nodes represent merged mRNA/protein components, sized and coloured by their dynamic importance (LL: light blue - dark red; LD: light lavender - dark purple, representing low to very high influence). Directed edges indicate regulatory influence, with colour and thickness categorised into quartiles based on interaction strength (light blue for weaker interactions to darker tones for stronger interactions). Each network was normalized independently within its own condition; therefore, node sizes and edge strengths represent relative influence within LL or LD separately and are not directly comparable across the two conditions.

The extended module comprising P51 and EL integrates into this core to stabilize phase transitions and maintain limit-cycle structure. P51 contributes to phase progression between morning and evening states, while EL modulates repression strength and supports oscillatory coherence. Cry shows high composite influence under LL, reflecting its strong role in constraining oscillatory dynamics through both period and phase portrait geometry. Composite edge weights further show that targeted degradation processes and selected translational steps influence waveform geometry without destabilizing rhythmicity, indicating a layered control structure. In contrast, photoreceptors PhyA and PhyB and downstream components such as COP1, PIFs, and GZ occupy peripheral positions with lower composite influence, consistent with their modulatory rather than rhythm-generating roles under LL.

Together, the integrated LL network depicts a compact transcription–degradation core that sustains autonomous oscillations, surrounded by light-input and translational modules that adjust amplitude, phase, and stability without redefining clock pace. This hierarchical organisation explains how the *Arabidopsis* circadian regulatory system maintains robust timing under constant illumination while preserving flexibility through distributed peripheral regulation.

Comparison of the LD and LL composite networks further clarifies how environmental light conditions reshape the distribution of regulatory influence within the clock (Figure 5A, B). Under LD cycles, influence spreads more broadly across network but redistribution differs from LL in several specific ways. P51 emerges as top hub in LD due to strong composite weights on P51 synthesis, translation and mRNA degradation (v3, p3, k3), reflecting its active role in light signal transmission under cycling conditions. Influence of Cry decreases indicating that Cry-dependent kinetic control is attenuated when the clock is externally driven. In contrast, under constant light the network becomes more internally consolidated: regulatory weight shifts toward the transcriptional core, particularly the CL-P97 module, which assumes stronger hub dominance and tighter connectivity with EL and Cry. Peripheral light-signalling components such as PhyA, COP1, and PIFs show reduced influence in LL, consistent with diminished entrainment input when external light transitions are absent. Together, these differences indicate that LD conditions distribute control towards P51, EL and light-input pathways including PhyA and PhyB, whereas LL condition emphasise the intrinsic transcription-degradation feedback structure, particularly the CL-P97 and Cry modules, that maintain autonomous circadian oscillations.

## 3. Discussion

The *Arabidopsis* circadian clock maintains precise daily timing despite fluctuating light environments and multilayer regulatory complexity. Understanding how transcriptional feedback, protein turnover, and light signalling integrate to sustain robust rhythms remains a central challenge in plant chronobiology(Harmer, 2009; Hsu & Harmer, 2014; McClung, 2006). A key motivation for developing improved computational model is the need to predict clock behaviour not only under controlled laboratory conditions but also under the complex, continuously varying light signals that plants experience in nature. The M1 model provides an improved and biologically grounded framework for dissecting this regulatory organisation and represents a step toward modelling circadian dynamics under natural conditions. By extending the Pay (2022) model to include additional light-dependent and transcriptional regulatory interactions, M1 resolves key discrepancies in gene expression dynamics that were not captured previously. Although the Pay model successfully simulated hypocotyl growth under extended photoperiods (6L:18D, 4L:20D, 2L:22D), it showed amplitude and, phase deviations relative to curated expression profiles when simulated under natural light conditions at equinoxes and solstices. Such deviation can disrupt coordination between internal rhythms and environmental light cycles, impairing photosynthetic efficiency, growth timing, and stress responses(Dodd et al., 2005; Greenham & McClung, 2015; Yerushalmi & Green, 2009). In contrast, M1 showed slightly improved correspondence to curated expression values under natural light conditions and modestly but consistently lower mean absolute error (MAE) and mean square error (MSE) relative to Pay model across all core components indicating modest improvement in predictive accuracy (Figure S2). These residual discrepancies highlight the inherent challenges of modelling circadian responses under natural conditions and point to components of the entrainment system that are not yet captured in the current framework. Further incorporation of the temperature entrainment, a well-established modulator of circadian phase and amplitude in plants, is expected to improve model prediction under natural conditions.

These improvements arise from incorporation of the GI/ZTL (GZ) module and additional regulatory links connecting COP1 to GZ, GZ to PRR5/TOC1 (P51), and CRY to PIF. These extensions strengthen coupling between transcriptional feedback and regulated protein stability, particularly within the evening and light-input modules. The added pathways regulate ubiquitin-mediated proteolysis of key clock proteins and modulate light-dependent transcriptional repression, processes known to shape circadian amplitude and phase stability(Kim et al., 2007; Más et al., 2000; Pedmale et al., 2016; Pokhilko et al., 2012). Notably, the GZ module parameters showed negligible period sensitivity and minimal phase portrait influence under both LL and LD conditions, consistent with a modulatory rather than rhythm-generating role. The improvement in predictive accuracy conferred by GZ incorporation therefore arises primarily through adjustment of amplitude and phase relationships in the evening loops rather than through direct changes in free-running period. This observation supports the view that GZ act as an interface between light-input pathways and the transcriptional core. It also explains why GZ knockout is classified as Class III despite its experimentally established importance for clock protein stability(Cha et al., 2017; Kiba et al., 2007; Más et al., 2003). Under natural conditions, where GI and ZTL activity is shaped by both light and temperature signals, incorporation of temperature entrainment is expected to further engage the amplitude and waveform control mechanisms identified in the phase portrait analysis and thereby improve model performance.

Knockout analysis under constant light highlights the central role of the reciprocal repression between CCA1/LHY (CL) and PRR9/PRR7 (P97) in maintaining circadian rhythmicity. Perturbations affecting CL production, turnover, or its inhibitory interactions with P97 consistently disrupted sustained oscillations, underscoring the importance of balanced synthesis, degradation, and mutual repression within this core feedback module. This mutual repression loop represents the established morning-afternoon feedback core of the *Arabidopsis* clock(Alabadí et al., 2001; Locke et al., 2006; Pokhilko et al., 2012). Period sensitivity analysis further demonstrated that synthesis and degradation rates within this core strongly constrain oscillation timing, emphasizing that turnover kinetics, in addition to transcriptional feedback, determine circadian pace. Control of protein stability is critical for matching internal rhythms to seasonal photoperiods and temperature cycles(Fujiwara et al., 2008; Gould et al., 2006). These findings are consistent with experimental studies showing that altered stability of CCA1 or PRR proteins modifies circadian period and amplitude(Green et al., 2002; Más et al., 2003).

Phase portrait analysis extended these results by revealing hierarchical control of oscillatory geometry. Parameters within the transcriptional repression core primarily influenced limit-cycle area, reflecting their role in sustaining oscillation amplitude and structural integrity. In contrast, selected degradation and translational processes altered eccentricity, indicating controlled modulation of waveform asymmetry without collapsing rhythmicity. Most peripheral light-input, degradation, and growth-output parameters produced only minor geometric changes, and the limit cycle remained closed across their tested ranges. Together, these results show that oscillatory stability is anchored by the CL-P97-P51-EL core, while translational and degradation processes refine waveform structure.

From a biological perspective, this organisation explains how plants maintain robust circadian timing under fluctuating environments. The transcription-degradation core functions as a self-sustained pacemaker under constant light, whereas light-regulated modules such as CRY-PIF and COP1-GZ provide entrainment flexibility and phase coordination under cycling conditions. Consistent with model results, CRY shows strong influence under constant light but reduced importance under LD conditions, reflecting decreased reliance on photoreceptor-driven control under external entrainment. This robustness ensures stable control of photosynthesis, hormone signalling, flowering time, and stress adaptation across variable day lengths(Dodd et al., 2005; Greenham & McClung, 2015; Song et al., 2010). The distributed and partially redundant light-input pathways observed in the model reflect experimentally documented resilience of the plant clock, where multiple regulatory layers buffer perturbations while preserving rhythmic precision(Michael et al., 2003; Webb et al., 2019). Such redundancy likely reflects evolutionary pressure to maintain reliable timekeeping under unpredictable light environments. Collectively, M1 captures key biological operating principles of the *Arabidopsis* circadian clock: a compact transcription-degradation core that generates rhythmicity, light-responsive modules that modulate phase and amplitude, and peripheral pathways that provide buffering and resilience. This hierarchical architecture reconciles structural necessity with quantitative flexibility, provides testable hypothesis for genetic perturbation and synthetic clock engineering, and offers a mechanistic framework for understanding circadian robustness in plants.

## 4. Methods

### 4.1 Model Framework

We developed an updated framework, M1 Model, by incorporating additional transcriptional and light-dependent regulatory interactions supported by experimental evidence. The model includes a new GI/ZTL (GZ) node representing the combined action of GIGANTEA and ZEITLUPE, which regulates protein stability and photoperiod sensing(Cha et al., 2017; Kiba et al., 2007; Más et al., 2003). Additional inhibitory interactions were introduced from GZ to P51 (PRR5/TOC1) to capture GI/ZTL mediated degradation control, from COP1 to GZ to simulate photoperiod-dependent modulation of GI/ZTL activity, and from CRY to PIF to reflect cryptochrome-mediated growth repression(Jang et al., 2015; Ma et al., 2016; Más et al., 2003). The new parameters were manually optimized. The complete system of ordinary differential equations (ODEs) and parameter lists are provided in the supplementary text and table S1.

The model was simulated under natural light conditions at equinoxes and solstices, and its predicted expression dynamics of CL (CCA1/LHY), P97 (PRR9/PRR7), P51 (PRR5/TOC1), and EL (ELF4/LUX) were analysed. The ODE system was solved numerically using the “odeint” function in Python (SciPy, odeint (LSODA-based) with a time step of 1 h and total duration of 240 hours(Virtanen et al., 2020). Model validation was carried out by comparing hypocotyl growth obtained from Pay et al., 2022 under combination of blue and red-light conditions with the three different photoperiods (6L:18D, 4L:20D and 2L:22D)(Pay et al., 2022). Further, we compared simulated expression profiles of CL (CCA1/LHY), P97 (PRR9/PRR7), P51 (PRR5/TOC1), and EL (ELF4/LUX) with curated expression data obtained from the Nagano et al., 2019 study(Nagano et al., 2019).

### 4.2 Knockout Analysis

To determine essential parameters required to sustain rhythmicity, each kinetic parameter was independently set to zero while all others were kept at reference values. The resulting time-series expression profiles for CL, P97, P51, and EL were analyzed to assess changes in period of oscillations. Further, the mean period change was calculated by calculating average change in period. Based on mean period change, parameters were classified into three classes: Class I, whose knockout caused complete loss of rhythmicity; Class II, which altered period but preserved rhythmicity; and Class III, which showed negligible impact.

### 4.3 Period Sensitivity Analysis

To quantify how variation in individual parameters affects the oscillation period, a systematic period sensitivity analysis was performed. Each kinetic parameter was varied independently across a 10-fold range (0.1x-10x baseline value) while keeping all others constant. The model was simulated for each value, and the oscillation period was determined from the time-series expression of four major components-CL, P97, P51, and EL. To avoid early transients, we excluded the first 240 time points from each simulation. We then normalised each time series by its maximum value and skipped signals with very low amplitude (amplitude range < 0.10). Peaks were detected using mild constraints (minimum height = 0.1, minimum prominence = 0.05) to ensure that only clear rhythmic maxima were selected. When at least three peaks were present, the period was calculated as the mean interval between successive peaks. The slope of period change with respect to parameter fold change was calculated to quantify sensitivity. Parameters were categorized as high, moderate, or low sensitivity based on their mean slope magnitude.

### 4.4 Phase Portrait Analysis

Phase portrait analysis was used to examine system dynamics and inter-component coupling. For each parameter, simulations were run for 480 h, and trajectories between selected component pairs CL-P97, CL-P51, CL-EL, P97-EL, P97-P51 and P51-EL were plotted in phase space after discarding the first 240 hours as transient behavior. Two geometric metrics were calculated from each trajectory: mean area, representing the oscillation amplitude or dynamic range, and eccentricity, representing the degree of phase asymmetry. Linear regression of each metric against the parameter fold change yielded slope-based indices describing how each parameter altered amplitude or shape.

### 4.5 Network Impact Analysis

Under constant light (LL) and light-dark cycle (LD), we constructed a composite weighted, directed gene-gene network by integrating the results of knockout analysis, period sensitivity analysis, and phase portrait analysis. Parameter-level results from the LL and LD combined table were merged with a curated parameter-to-edge mapping file specifying source, target, and interaction type. For each parameter, knockout class (Class I = 1.0, Class II = 0.5, Class III = 0.0) was multiplied by the normalized absolute knockout-induced period change to obtain a structural term (K_norm). Period sensitivity (P_norm), phase portrait area change (A_norm), and eccentricity change (E_norm) were normalized by their maximum absolute values within LL and LD, individually.

A composite edge weight was then defined as-

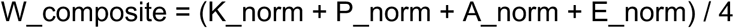

with equal weighting across metrics. This unified score captures structural necessity, period control, and waveform modulation in a single quantitative measure. A directed graph was constructed using W_composite as edge weight, and node influence was assessed using strength and weighted centrality measures. The resulting LL and LD composite networks therefore represents the integrated regulatory architecture under constant light and light-dark cycle, respectively, combining essentiality and dynamical impact within a single framework.

## Acknowledgement

SS acknowledges financial support by Indian Institute of Technology Gandhinagar and Ministry of Education, India. AS acknowledges support from DBT Ramalingaswami Re-entry Fellowship and Indian Institute of Technology Gandhinagar.

## Authors Contributions

Shashank Kumar Singh designed and implemented the mathematical framework, carried out simulations and computational analyses, interpreted the data, prepared the figures, and wrote the manuscript. Ashutosh Srivastava conceived and supervised the study, provided scientific guidance throughout the work, contributed to interpretation of the results, and revised the manuscript critically.

## Data Availability

All code used for model simulation, parameter analysis, and network construction in this study is available at GitHub: https://github.com/clocklabiitgn/M1_Plant_Circadian_Network_Architecture

The repository includes the implementation of the M1 circadian clock model and scripts used for knockout analysis, period sensitivity analysis, phase portrait analysis, and composite network construction under light-dark (LD) and constant light (LL) conditions.

