## Supplementary File for "Dissecting the Network Architecture of a Plant Circadian Clock Model: Identifying Key Regulatory Mechanisms and Essential Interactions"

### 1 Supplementary 1: Equations of the M1 model

The equations are similar to those of the Pay model 2022b. Equations 19 is added along with modifications in Equations 6 and 10 of P51 protein and PIF mRNA, respectively.

$$\frac{d[\mathrm{CLm}]}{dt} = (v_1 + L_a)\left(\frac{1}{1 + \left(\frac{[\mathrm{P97p}]}{K_1}\right)^2 + \left(\frac{[\mathrm{P51p}]}{K_2}\right)^2}\right) - \big(k_{1L}\Theta_{\mathrm{PhyA}} + k_{1D}(1 - \Theta_{\mathrm{PhyA}})\big)[\mathrm{CLm}] \tag{1}$$

$$\frac{d[\mathrm{CLp}]}{dt} = \big(p_1 + p_{1L}\Theta_{\mathrm{PhyA}}\big)[\mathrm{CLm}] - d_1[\mathrm{CLp}] \tag{2}$$

$$\frac{d[\mathrm{P97m}]}{dt} = (v_2 + L_b)\left(\frac{1}{1 + \left(\frac{[\mathrm{CLp}]}{K_3}\right)^2 + \left(\frac{[\mathrm{P51p}]}{K_4}\right)^2 + \left(\frac{[\mathrm{ELp}]}{K_5}\right)^2}\right) - k_2[\mathrm{P97m}] \tag{3}$$

$$\frac{d[\mathrm{P97p}]}{dt} = p_2[\mathrm{P97m}] - d_{2D}(1 - \Theta_{\mathrm{PhyA}})[\mathrm{P97p}] - d_{2L}\Theta_{\mathrm{PhyA}}[\mathrm{P97p}] \tag{4}$$

$$\frac{d[\mathrm{P51m}]}{dt} = \left(\frac{v_3}{1 + \left(\frac{[\mathrm{CLp}]}{K_6}\right)^2 + \left(\frac{[\mathrm{P51p}]}{K_7}\right)^2}\right) - k_3[\mathrm{P51m}] \tag{5}$$

$$\frac{d[\mathrm{P51p}]}{dt} = p_3[\mathrm{P51m}] - d_{3D}(1 - \Theta_{\mathrm{PhyA}})[\mathrm{P51p}] - d_{3L}\Theta_{\mathrm{PhyA}}[\mathrm{P51p}] - \frac{d_{p6}[\mathrm{GZp}][\mathrm{P51p}]}{G_{kp} + [\mathrm{P51p}]} \tag{6}$$

$$\frac{d[\mathrm{ELm}]}{dt} = \left(\frac{v_4\Theta_{\mathrm{PhyA}}}{1 + \left(\frac{[\mathrm{CLp}]}{K_8}\right)^2 + \left(\frac{[\mathrm{P51p}]}{K_9}\right)^2 + \left(\frac{[\mathrm{ELp}]}{K_{10}}\right)^2}\right) - k_4[\mathrm{ELm}] \tag{7}$$

$$\begin{aligned}\frac{d[\mathrm{ELp}]}{dt} = p_4[\mathrm{ELm}] - \Bigg(d_{e1} &+ \frac{d_{e2}[\mathrm{COP1}] + d_{e3}[\mathrm{COP1:PhyA}]}{[\mathrm{COP1}] + [\mathrm{COP1:PhyA}] + [\mathrm{COP1:PhyB}] + [\mathrm{COP1:Cry1}]} \\ &+ \frac{d_{e4}[\mathrm{COP1:PhyB}] + d_{e5}[\mathrm{COP1:Cry1}]}{[\mathrm{COP1}] + [\mathrm{COP1:PhyA}] + [\mathrm{COP1:PhyB}] + [\mathrm{COP1:Cry1}]}\Bigg)[\mathrm{ELp}]\end{aligned} \tag{8}$$

$$\begin{aligned}\frac{d[\mathrm{PhyA}]}{dt} &= (1 - \Theta_{\mathrm{PhyA}})A_{p3} - \frac{A_{m7}[\mathrm{PhyA}]}{A_{k7} + [\mathrm{PhyA}]} \\ &- q_2\Theta_{\mathrm{PhyA}}[\mathrm{PhyA}] - k_{mpac}\Theta_{\mathrm{PhyA}}[\mathrm{PhyA}][\mathrm{COP1}] \\ &+ k_d[\mathrm{COP1:PhyA}]\end{aligned} \tag{9}$$

$$\frac{d[\mathrm{PIFm}]}{dt} = \frac{v_5}{1 + \left(\frac{[\mathrm{ELp}]}{K_{11}}\right)^2 + \left(\frac{[\mathrm{Cry1}]}{K_{14}}\right)^2} - k_5[\mathrm{PIFm}] \tag{10}$$

$$\frac{d[\mathrm{PIFp}]}{dt} = p_5[\mathrm{PIFm}] - d_{5D}(1 - \Theta_{\mathrm{PhyA}})[\mathrm{PIFp}] - d_{5L}\Theta_{\mathrm{PhyA}}[\mathrm{PIFp}] \tag{11}$$

$$\frac{d[\mathrm{HYPp}]}{dt} = g_1 + \frac{g_2[\mathrm{PIFp}]^2}{K_{12}^2 + [\mathrm{PIFp}]^2} \tag{12}$$

$$\begin{aligned}\frac{d[\mathrm{PhyB}]}{dt} &= B_{p4} - \frac{B_{m8}[\mathrm{PhyB}]}{B_{k8} + [\mathrm{PhyB}]} \\ &- k_{mpbc}\Theta_{\mathrm{PhyB}}[\mathrm{PhyB}][\mathrm{COP1}] + k_d[\mathrm{COP1} : \mathrm{PhyB}]\end{aligned} \tag{13}$$

$$\begin{aligned}\frac{d[\mathrm{Cry1}]}{dt} &= C_{p5} - \frac{C_{m9}[\mathrm{Cry1}]}{C_{k9} + [\mathrm{Cry1}]} \\ &- k_{mcc}\Theta_{\mathrm{Cry1}}[\mathrm{Cry1}][\mathrm{COP1}] + k_d[\mathrm{COP1} : \mathrm{Cry1}]\end{aligned} \tag{14}$$

$$\begin{aligned}\frac{d[\mathrm{COP1}]}{dt} &= -k_{mpac}\Theta_{\mathrm{PhyA}}[\mathrm{PhyA}][\mathrm{COP1}] + k_d[\mathrm{COP1} : \mathrm{PhyA}] \\ &- k_{mpbc}\Theta_{\mathrm{PhyB}}[\mathrm{PhyB}][\mathrm{COP1}] + k_d[\mathrm{COP1} : \mathrm{PhyB}] \\ &- k_{mcc}\Theta_{\mathrm{Cry1}}[\mathrm{Cry1}][\mathrm{COP1}] + k_d[\mathrm{COP1} : \mathrm{Cry1}] \\ &+ \frac{A_{m7}[\mathrm{COP1} : \mathrm{PhyA}]}{A_{k7} + [\mathrm{COP1} : \mathrm{PhyA}]} + q_2\Theta_{\mathrm{PhyA}}[\mathrm{COP1} : \mathrm{PhyA}] \\ &+ \frac{B_{m8}[\mathrm{COP1} : \mathrm{PhyB}]}{B_{k8} + [\mathrm{COP1} : \mathrm{PhyB}]} + \frac{C_{m9}[\mathrm{COP1} : \mathrm{Cry1}]}{C_{k9} + [\mathrm{COP1} : \mathrm{Cry1}]}\end{aligned} \tag{15}$$

$$\begin{aligned}\frac{d[\mathrm{COP1} : \mathrm{PhyA}]}{dt} &= k_{mpac}\Theta_{\mathrm{PhyA}}[\mathrm{PhyA}][\mathrm{COP1}] - k_d[\mathrm{COP1} : \mathrm{PhyA}] \\ &- \frac{A_{m7}[\mathrm{COP1} : \mathrm{PhyA}]}{A_{k7} + [\mathrm{COP1} : \mathrm{PhyA}]} - q_2\Theta_{\mathrm{PhyA}}[\mathrm{COP1} : \mathrm{PhyA}]\end{aligned} \tag{16}$$

$$\begin{aligned}\frac{d[\mathrm{COP1} : \mathrm{PhyB}]}{dt} &= k_{mpbc}\Theta_{\mathrm{PhyB}}[\mathrm{PhyB}][\mathrm{COP1}] - k_d[\mathrm{COP1} : \mathrm{PhyB}] \\ &- \frac{B_{m8}[\mathrm{COP1} : \mathrm{PhyB}]}{B_{k8} + [\mathrm{COP1} : \mathrm{PhyB}]}\end{aligned} \tag{17}$$

$$\begin{aligned}\frac{d[\mathrm{COP1} : \mathrm{Cry1}]}{dt} &= k_{mcc}\Theta_{\mathrm{Cry1}}[\mathrm{Cry1}][\mathrm{COP1}] - k_d[\mathrm{COP1} : \mathrm{Cry1}] \\ &- \frac{C_{m9}[\mathrm{COP1} : \mathrm{Cry1}]}{C_{k9} + [\mathrm{COP1} : \mathrm{Cry1}]}\end{aligned} \tag{18}$$

$$\begin{aligned}\frac{d[\mathrm{GZp}]}{dt} &= G_{p6} - \left(d_{g1} + \frac{d_{g2}[\mathrm{COP1}] + d_{g3}[\mathrm{COP1} : \mathrm{PhyA}]}{[\mathrm{COP1}] + [\mathrm{COP1} : \mathrm{PhyA}] + [\mathrm{COP1} : \mathrm{PhyB}] + [\mathrm{COP1} : \mathrm{Cry1}]}\right. \\ &\left. + \frac{d_{g4}[\mathrm{COP1} : \mathrm{PhyB}] + d_{g5}[\mathrm{COP1} : \mathrm{Cry1}]}{[\mathrm{COP1}] + [\mathrm{COP1} : \mathrm{PhyA}] + [\mathrm{COP1} : \mathrm{PhyB}] + [\mathrm{COP1} : \mathrm{Cry1}]}\right)[\mathrm{GZp}]\end{aligned} \tag{19}$$

**Light activation term:**

$$L_u = \Big(q_{1u}\big([\text{PhyA}]\big)\Theta_{\text{PhyA}}\Big) + \Big(q_{3u}\big([\text{PhyB}]\big)\log_{10}(\eta_1 I_{\text{red}}+1)\Theta_{\text{PhyB}}\Big)$$

$$+\Big(q_{4u}\big([\text{Cry1}]\big)\log_{10}(\eta_2 I_{\text{blue}}+1)\Theta_{\text{Cry1}}\Big)$$

Here, $u = $ (a or b) for CL and P97, respectively.

For $\Theta_{PhyA}, \Theta_{PhyB}$ and $\Theta_{Cry1}$ values, the following conditions are followed-

$$\Theta_{PhyA} = \begin{cases} 1, & I_{red} \text{ or } I_{blue} \neq 0, \\ 0, & \text{otherwise} \end{cases}$$

$$\Theta_{PhyB} = \begin{cases} 1, & I_{red} \neq 0, \\ 0, & \text{otherwise} \end{cases}$$

$$\Theta_{Cry1} = \begin{cases} 1, & I_{blue} \neq 0, \\ 0, & \text{otherwise} \end{cases}$$

| Table S1- Model parameters used in the M1 circadian clock model. | | | | |
|---|---|---|---|---|
| **S.No** | **Description** | **Symbol** | **Value** | **Unit** |
| 1 | CL synthesis | **v1** | 4.8318 | $nM \cdot h^{-1}$ |
| 2 | CL light-induced synthesis through PhyA | **q1a** | 1.4266 | $nM \cdot h^{-1}$ |
| 3 | CL light-induced synthesis through PhyB | **q3a** | 8.9432 | $nM \cdot h^{-1}$ |
| 4 | Normalization of red light intensity | **eta1** | 0.03 | - |
| 5 | CL light-induced synthesis through Cry | **q4a** | 5.9277 | $nM \cdot h^{-1}$ |
| 6 | Normalization of blue light intensity | **eta2** | 0.0215 | - |
| 7 | Inhibition: CL by P97 | **K1** | 0.1943 | nM |
| 8 | Inhibition: CL by P51 | **K2** | 1.6138 | nM |
| 9 | CL mRNA degradation (light) | **k1L** | 0.2866 | $h^{-1}$ |
| 10 | CL mRNA degradation (dark) | **k1D** | 0.213 | $h^{-1}$ |
| 11 | CL translation | **p1** | 0.8672 | $h^{-1}$ |
| 12 | CL light-induced translation | **p1L** | 0.2378 | $h^{-1}$ |
| 13 | CL degradation | **d1** | 0.7843 | $h^{-1}$ |
| 14 | P97 light-induced synthesis through PhyA | **q1b** | 3.575 | $nM \cdot h^{-1}$ |
| 15 | P97 light-induced synthesis through PhyB | **q3b** | 5.5899 | $nM \cdot h^{-1}$ |
| 16 | P97 light-induced synthesis through Cry | **q4b** | 8.954 | $nM \cdot h^{-1}$ |
| 17 | P97 synthesis | **v2** | 1.6822 | $nM \cdot h^{-1}$ |
| 18 | Inhibition: P97 by CL | **K3** | 2.2275 | nM |
| 19 | Inhibition: P97 by P51 | **K4** | 0.4 | nM |
| 20 | Inhibition: P97 by EL | **K5** | 0.37 | nM |
| 21 | P97 mRNA degradation | **k2** | 0.35 | $h^{-1}$ |
| 22 | P97 translation | **p2** | 0.7858 | $h^{-1}$ |
| 23 | P97 degradation (dark) | **d2D** | 0.3712 | $h^{-1}$ |
| 24 | P97 degradation (light) | **d2L** | 0.2917 | $h^{-1}$ |
| 25 | P51 synthesis | **v3** | 1.113 | $nM \cdot h^{-1}$ |
| 26 | Inhibition: P51 by CL | **K6** | 0.4944 | nM |
| 27 | Inhibition: P51 by itself | **K7** | 2.4087 | nM |
| 28 | P51 mRNA degradation | **k3** | 0.5819 | $h^{-1}$ |
| 29 | P51 translation | **p3** | 0.6142 | $h^{-1}$ |
| 30 | P51 degradation (dark) | **d3D** | 0.5026 | $h^{-1}$ |
| 31 | P51 degradation (light) | **d3L** | 0.5431 | $h^{-1}$ |
| 32 | EL synthesis | **v4** | 2.5012 | $nM \cdot h^{-1}$ |
| 33 | Inhibition: EL by CL | **K8** | 0.3262 | nM |
| 34 | Inhibition: EL by P51 | **K9** | 1.7974 | nM |
| 35 | Inhibition: EL by EL | **K10** | 1.1889 | nM |
| 36 | EL mRNA degradation | **k4** | 0.925 | $h^{-1}$ |
| 37 | EL translation | **p4** | 1.126 | $h^{-1}$ |
| 38 | EL degradation | **de1** | 0.0022 | $h^{-1}$ |
| 39 | EL degradation (COP1) | **de2** | 0.4741 | $h^{-1}$ |
| 40 | EL degradation (COP1: PhyA) | **de3** | 0.3765 | $h^{-1}$ |
| 41 | EL degradation (COP1: PhyB) | **de4** | 0.398 | $h^{-1}$ |
| 42 | EL degradation (COP1: Cry) | **de5** | 0.0003 | $h^{-1}$ |
| 43 | PhyA translation | **Ap3** | 0.3868 | $h^{-1}$ |
| 44 | PhyA degradation | **Am7** | 0.5503 | $h^{-1}$ |
| 45 | Michaelis constant of PhyA degradation | **Ak7** | 1.125 | nM |
| 46 | Rate constant of light-independent degradation | **q2** | 0.5767 | $h^{-1}$ |
| 47 | Binding rate of COP1:PhyA | **kmpac** | 137 | $nM^{-1} \cdot h^{-1}$ |

| | | | | |
|---|---|---|---|---|
| 48 | Dissociation rate | **kd** | 7 | $h^{-1}$ |
| 49 | PIF synthesis | **v5** | 0.1129 | $nM \cdot h^{-1}$ |
| 50 | Inhibition: PIF by EL | **K11** | 0.3322 | nM |
| 51 | PIF mRNA degradation | **k5** | 0.1591 | $h^{-1}$ |
| 52 | PIF translation | **p5** | 0.5293 | $h^{-1}$ |
| 53 | PIF protein degradation (dark) | **d5D** | 0.4404 | $h^{-1}$ |
| 54 | PIF protein degradation (light) | **d5L** | 5.0712 | $h^{-1}$ |
| 55 | Baseline hypocotyl growth | **g1** | 0.001 | $mm \cdot h^{-1}$ |
| 56 | PIF-induced hypocotyl growth | **g2** | 0.18 | $mm \cdot h^{-1}$ |
| 57 | Activation: growth by PIF | **K12** | 0.86 | nM |
| 58 | PhyB translation | **Bp4** | 0.4147 | $h^{-1}$ |
| 59 | PhyB degradation | **Bm8** | 0.7728 | $h^{-1}$ |
| 60 | Michaelis constant of PhyB degradation | **Bk8** | 0.1732 | nM |
| 61 | Binding rate of COP1:PhyB | **kmpbc** | 7162 | $nM^{-1} \cdot h^{-1}$ |
| 62 | Cry translation | **Cp5** | 0.4567 | $h^{-1}$ |
| 63 | Cry degradation | **Cm9** | 0.867 | $h^{-1}$ |
| 64 | Michaelis constant of Cry degradation | **Ck9** | 0.3237 | nM |
| 65 | Binding rate of COP1:Cry | **kmcc** | 13406 | $nM^{-1} \cdot h^{-1}$ |
| 66 | Inhibition: PIF by Cry | **K14** | 1.5 | nM |
| 67 | GZ translation | **Gp6** | 0.0001 | $nM \cdot h^{-1}$ |
| 68 | GZ degradation | **dg1** | 0.01 | $h^{-1}$ |
| 69 | GZ degradation (COP1) | **dg2** | 1.280202 | $h^{-1}$ |
| 70 | GZ degradation (COP1: PhyA) | **dg3** | 0.01 | $h^{-1}$ |
| 71 | GZ degradation (COP1: PhyB) | **dg4** | 1.750462 | $h^{-1}$ |
| 72 | GZ degradation (COP1: Cry) | **dg5** | 1.067661 | $h^{-1}$ |
| 73 | P51 degradation (GZ) | **dp6** | 0.01 | $h^{-1}$ |
| 74 | Michaelis constant of P51 degradation by GZ | **Gkp** | 1.185527 | nM |

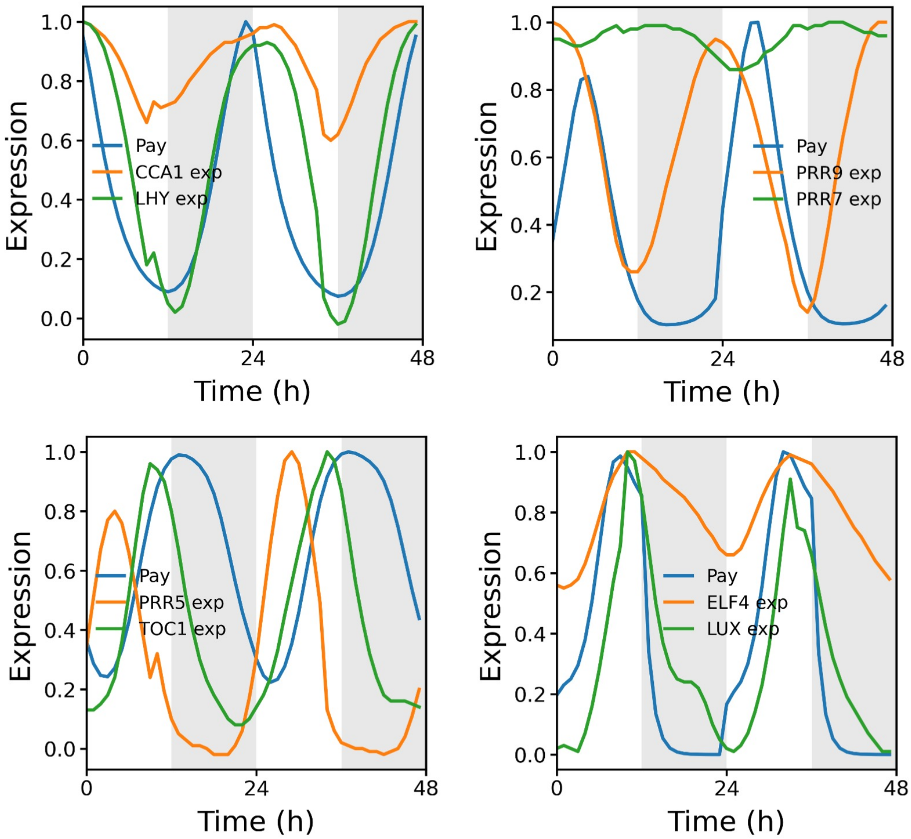


**Figure S1 –** Expression of core components of Pay 2022 model under natural condition on spring equinox with experimental data extracted from the Nagano et al., 2019.

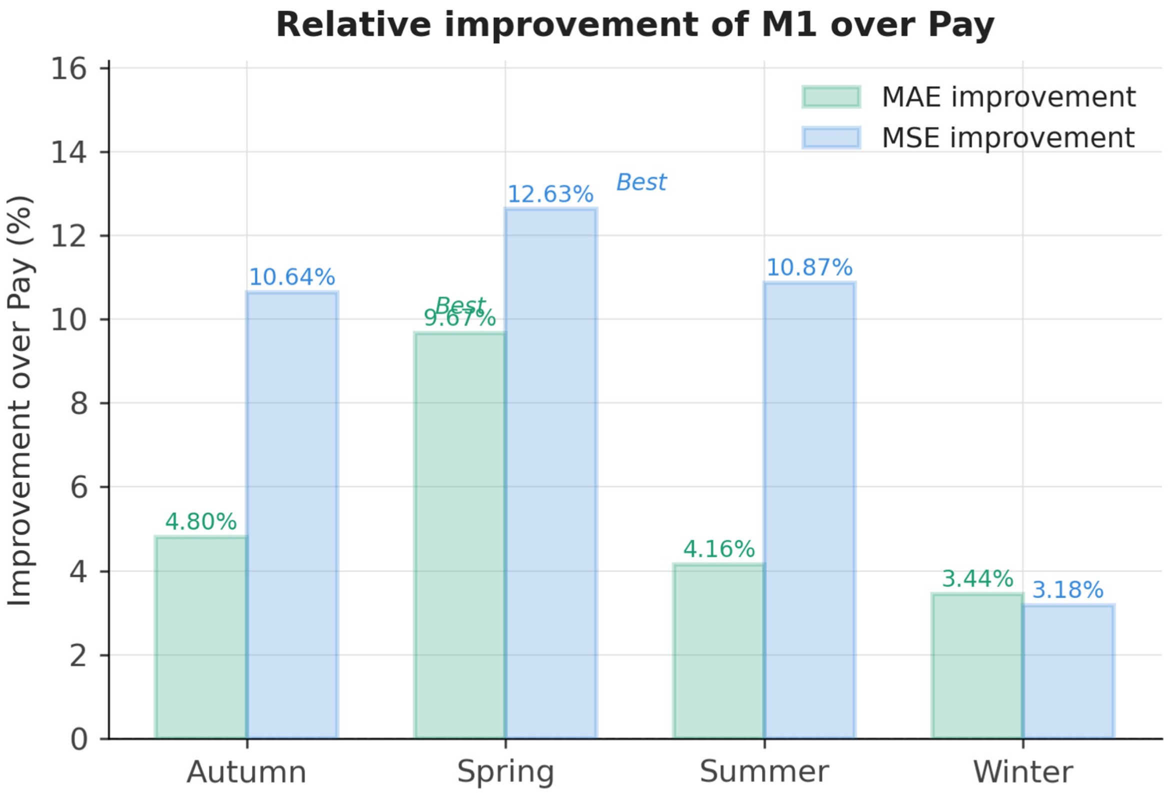


**Figure S2 –** Relative improvement in predictions of M1 model over Pay model predictions in terms of Mean Absolute Error (MAE) and Mean Square Error (MSE) across all 4 seasons.
